# Multi-state design of kinetically-controlled RNA aptamer ribosensors

**DOI:** 10.1101/213538

**Authors:** Cassandra R. Burke, David Sparkman-Yager, James M. Carothers

**Affiliations:** Department of Chemical Engineering Department of Bioengineering Molecular Engineering & Sciences Institute Center for Synthetic Biology University of Washington Seattle, WA 98195 United States

## Abstract

Metabolite-responsive RNA regulators with kinetically-controlled responses are widespread in nature. By comparison, very limited success has been achieved creating kinetic control mechanisms for synthetic RNA aptamer devices. Here, we show that kinetically-controlled RNA aptamer ribosensors can be engineered using a novel approach for multi-state, co-transcriptional folding design. The design approach was developed through investigation of 29 candidate *p*-aminophenylalanine-responsive ribosensors. We show that ribosensors can be transcribed *in situ* and used to analyze metabolic production directly from engineered microbial cultures, establishing a new class of cell-free biosensors. We found that kinetically-controlled ribosensors exhibited 5-10 fold greater ligand sensitivity than a thermodynamically-controlled device. And, we further demonstrated that a second aptamer, promiscuous for aromatic amino acid binding, could be assembled into kinetic ribosensors with 45-fold improvements in ligand selectivity. These results have broad implications for engineering RNA aptamer devices and overcoming thermodynamic constraints on molecular recognition through the design of kinetically-controlled responses.

## INTRODUCTION

Metabolite-responsive RNA regulators that respond to changing conditions through molecular interactions are widespread in biology^1^. In many of these systems, kinetic control mechanisms coordinate co-transcriptional RNA folding with metabolite binding and enable responses that are highly-sensitive and highly-selective to target ligands^2–6^. Although synthetic riboswitches exhibiting kinetic control have been identified by chance^7,8^, it has not been possible to intentionally engineer kinetically-controlled RNA aptamer devices^9^. Consequently, kinetic control mechanisms that could otherwise be exploited to overcome functional limits imposed by the thermodynamics of molecular recognition have remained beyond reach^10–13^.

Kinetic control is characterized by irreversibility and rate-driven outcomes that do not depend on the thermodynamic stabilities of the final products^14^. A key feature of kinetically-controlled RNA regulators is the co-transcriptional binding window, a period of time for the RNA to interact with the metabolite, that closes with the transition to an irreversible non-signaling state^5^. Changing the duration of the binding window provides a route for tuning the response^15^. Lengthening the binding window increases aptamer-ligand association, increasing the sensitivity of the RNA to lower metabolite concentrations. For example, the *E. coli* thiamine pyrophosphate (TPP) riboswitches have binding windows as large as 12 seconds, conferring 10-fold increases in ligand sensitivity^5^. By a similar rationale, binding windows that allow molecular discrimination on the basis of RNA-ligand association rates can dramatically increase the selectivity for a target even when the equilibrium binding affinities among cognate molecules are very similar^6^. Therefore, developing the ability to reliably access kinetic control would create unique capabilities for engineering ligand sensitivity and selectivity that are not possible with thermodynamic control^13^.

Here, we show that kinetically-controlled RNA aptamer ‘ribosensors’ can be engineered using a novel approach for multi-state, co-transcriptional RNA folding design. *In vitro* selected RNA aptamers are coupled through a timer domain to a toehold-mediated strand displacement (TMSD) actuator. In this architecture, co-transcriptional ligand-binding generates fluorescent outputs from DNA gates through TMSD (Fig. 1a-b)^16^. Using an RNA aptamer that binds an industrially- and medically-relevant aromatic amino acid (*p*-amino-phenylalanine, *p*-AF)(**Supplementary Fig. S1**) as a starting point^17–19^, we engineered kinetically-controlled ribosensors that can be transcribed *in situ* and produce fluorescent outputs measured in realtime without further purification or preparation. We validated the kinetic control mechanism by selectively inhibiting RNA polymerase and measuring the impact on ribosensor output. Through the intentional design of a thermodynamically-controlled device, we show how the underlying design parameters give rise to kinetic control and observe that kinetically-controlled devices exhibit significantly more sensitivity toward the target ligand.

**Figure 1.**
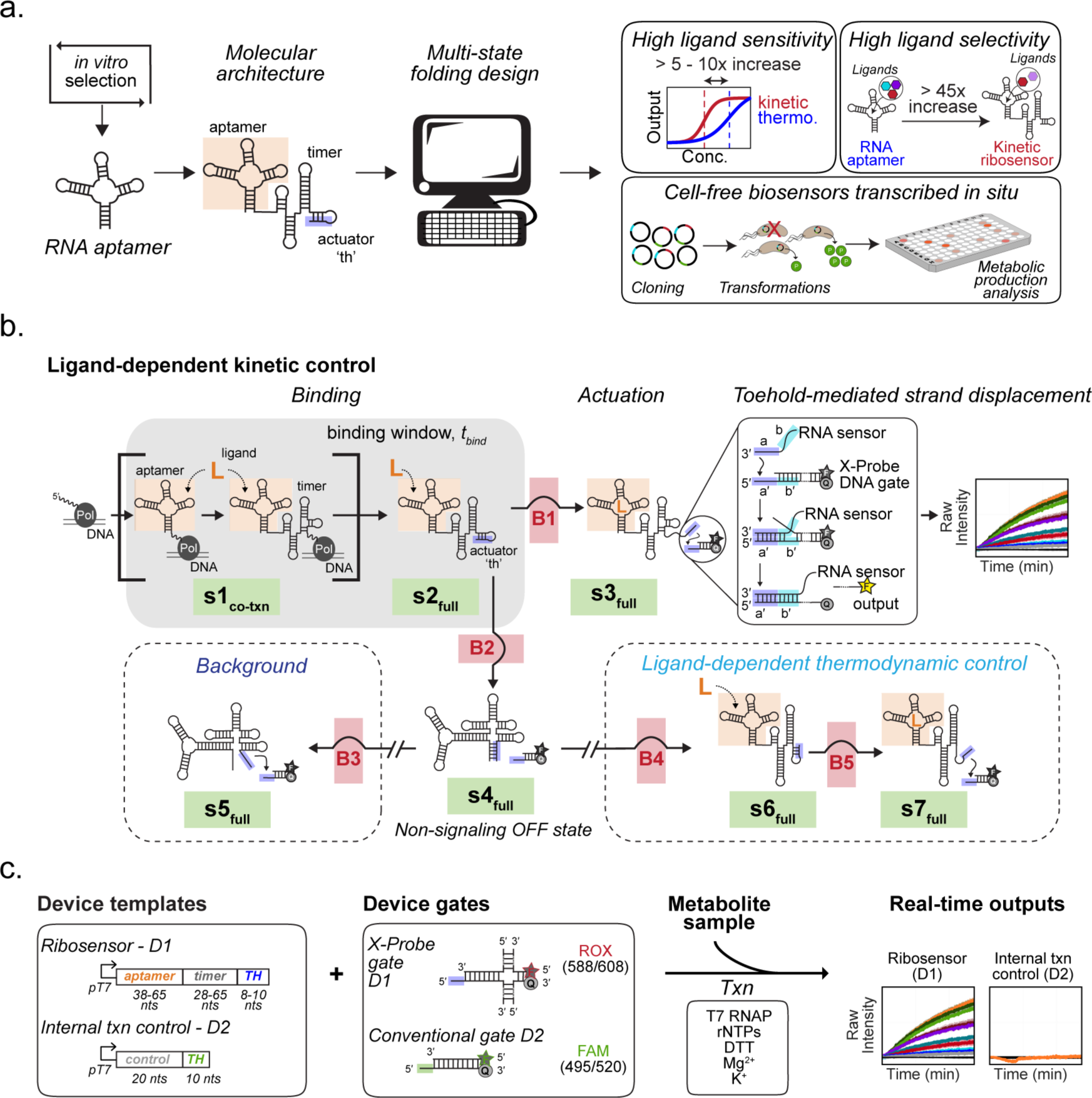
Kinetically-controlled ribosensors. (a) Kinetically-controlled ribosensors are engineered from *in vitro* selected RNA aptamers through multi-state co-transcriptional folding design. In this molecular architecture, a ‘timer’ domain couples the aptamer to a toehold-mediated strand displacement (TMSD) actuator (‘th’). Kinetically-controlled ribosensors can circumvent thermodynamic constraints on molecular recognition and are functional as cell-free biosensors for metabolic production analysis. (b) Secondary structure states (green) and free energy barriers (red) are designed to frame a binding window (given by *t*_*bind*_), during which the RNA can associate with the ligand. Ligand-dependent, kinetically-controlled responses generate FRET-based fluorescent outputs when the toehold actuator (th) interacts with the DNA gate through TMSD. (c) Ribosensors are transcribed *in situ*, alongside internal transcription controls, in the presence of fluorescently-labeled TMSD gates and the metabolite sample. Ligand-dependent fluorescent outputs are analyzed in real-time.

By quantifying metabolic production titers directly from engineered *E. coli* cell culture media, we establish these devices as a new class of low-cost biosensors with immediate, practical utility. And, we further demonstrate the generality of the *p*-AF ribosensor-derived design parameters by assembling a second aptamer, promiscuous for aromatic amino acid binding^20,21^, into a highly-selective ribosensor. Taken together, these results show that functional ribosensors can be designed for real-world applications and highlight the potential for harnessing kinetic control mechanisms to engineer highly-sensitive and highly-selective RNA aptamer devices.

## RESULTS

### Multi-state co-transcriptional design of functional devices

We began by formulating a molecular architecture to permit the assembly of *in vitro* selected RNA aptamers into functional ribosensors with ligand-dependent, kinetically-controlled responses (Fig. 1). Briefly, the design goal is to create devices with aptamers that can interact with the ligand within a binding window (defined by *t*_*bind*_) before the RNA folds into an irreversible non-signaling OFF state. Aptamer-ligand association occurring within the binding window results in actuation through an RNA structure state that initiates a TMSD reaction, generating concentration-dependent fluorescent outputs. At a minimum, we needed an approach for generating RNA sequences that fold along reliable co-transcriptional pathways into distinct, but ligand-dependent, structures for binding and actuation.

Secondary structure design based on minimal free energy (MFE) predictions can identify sequences capable of folding into multiple states^22,23^. The structure states themselves are determined by the energy landscape of the ensemble, consisting of free energy basins separated by free energy barriers^8^. We hypothesized that if the energy barriers separating structures along co-transcriptional folding trajectories could be predicted^9,24^, then sequences with targeted rates of transition from one state to another could be identified^25^. In this way, binding windows for aptamer-ligand association could be created through secondary structure design, providing an avenue to engineer kinetically-controlled responses.

Using these principles, we developed a novel approach for engineering kinetically-controlled ribosensors using multi-state, co-transcriptional secondary structure folding design (Fig. 2a). Secondary structure states and free energy barriers are designed to keep the RNA from folding into the OFF state too quickly, which would prematurely close the binding window and prevent the aptamer from interacting with the ligand. The binding window is further framed by structure states and free energy barriers preventing the RNA from folding directly into the actuation state, minimizing the output except when ligand binding has occurred (Fig. 1b). In this design, the outputs are dictated by differences in free energies of the secondary structures and the intervening barriers.

**Figure 2.**
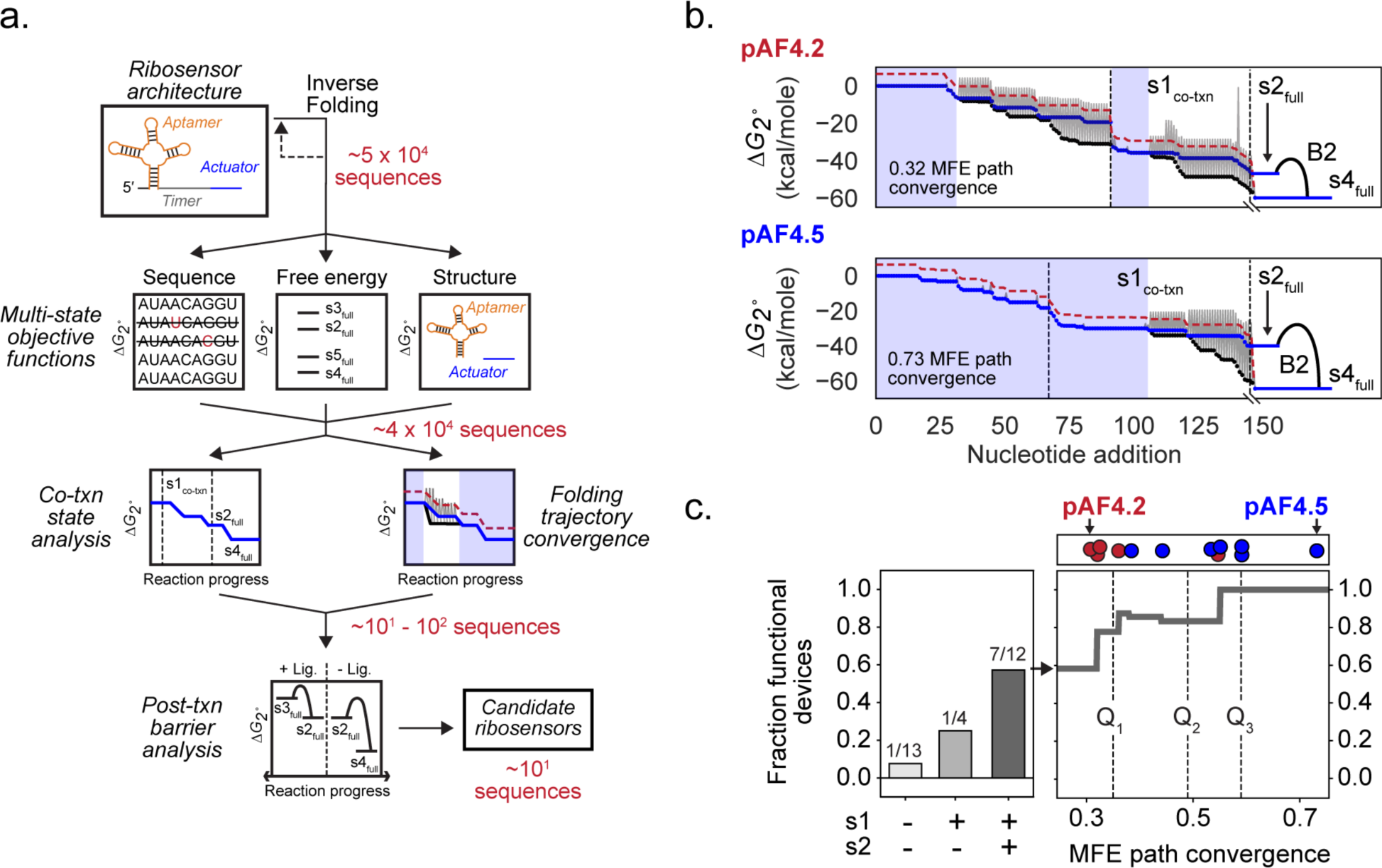
Multi-state co-transcriptional RNA folding design. (a) First, inverse folding is used to generate sequences satisfying multi-state objective functions for the full-length sequence, secondary structure free energies and targeted secondary structure states. Next, MFEpath is used to evaluate the accessibility of the targeted structure states along the predicted co-transcriptional folding trajectories, and the degree of convergence with the pathway defined by the elongating substructure minimal free energies (MFEs). Finally, direct folding pathways are analyzed to calculate the rates of transition from the final *MFEpath*-predicted post-transcriptional structure to the non-signaling OFF state. (b) MFE path convergence (blue shading) is plotted for two candidate *p*-AF ribosensors as the number of elongation steps along the co-transcriptional folding pathway where the predicted secondary structure (blue line) matches the target MFE substructure (black line). The free energy barriers between each step (gray lines) and the transition thresholds are shown (red dashed line). (c) Initially, 29 candidate *p*-AF ribosensors were designed and tested to investigate the design approach. The fraction of functional *p*-AF-responsive ribosensors is plotted against the predicted accessibility of the s1_co-txn_ and s2_full_ structure states (light to dark gray bars) and MFE path convergence; functional (blue) and non-functional devices (red) are indicated.

We created a hierarchical computational approach to generate ribosensor sequences meeting targeted design specifications (Fig. 2a). First, secondary structure objective functions for the full-length sequences are fed into an inverse folding genetic algorithm (**Supplementary Fig. S2**)^26^. Next, the accessibility of targeted co-transcriptional structures is evaluated using *MFEpath*. *MFEpath* simulates co-transcriptional folding trajectories as a series of binary decisions by comparing the rate of folding into the MFE substructure at each elongation step with the rate of nucleotide addition (Fig. 2b, **Supplementary Fig. S3** and **Supplementary Tables S1, S2**)^25,27–29^. Finally, direct folding pathways are analyzed to calculate the rates of transition from the final, *MFEpath*-predicted, post-transcriptional structure (*s2*_*full*_) to either the ligand-bound actuation state (*s3*_*full*_) or the non-signaling OFF state (*s4*_*full*_) (**Supplementary Table S3**)^29^.

To evaluate the design approach, we generated ~10^4^ sequences through inverse folding design and then used our computational pipeline (Fig. 2a) to reduce the set to 29 candidate ribosensors comprised of a 65 nucleotide (nt) *p*-aminophenylalanine (*p*-AF)-binding RNA aptamer^17,18^, a 65 nt timer and a 10 nt toehold actuator (Fig. 1c, **Supplementary Tables S4, S5**). The set of candidate ribosensors contained sequences with differences in the multi-state secondary structure free energies and barriers (**Supplementary Tables S1 - S3**). There were also wide variations in the predicted accessibility of the targeted structure states along the simulated co-transcriptional folding trajectories (Fig. 2c). In addition, there were large differences in the degree to which the RNAs were expected to fold, or converge, on the pathway defined by the elongating substructure MFEs (Figs. 2b, 2c and **Supplementary Table S2**). By optimizing reaction conditions for compatibility with both T7 RNA polymerase and TMSD^16^, the ribosensors can be transcribed *in situ* and produce ligand-dependent fluorescent outputs analyzed in real-time (Fig. 1c). In these assays, 9 of the resulting sequences exhibited *p*-AF-responsive outputs, including 7 of 12 sequences with ligand-binding states (*s1*_*co-txn*_ and *s2*_*full*_) predicted to be accessible along the co-transcriptional folding trajectory (Fig. 2c, **Supplementary Table S2**). In contrast, only 1 of 13 devices without these ligand-binding states in the simulated folding trajectory gave *p*-AF-dependent responses. Notably, for the 7 devices satisfying the co-transcriptional structure state criteria, there is excellent correspondence between function and the degree of MFE path convergence (Figs. 2b, 2c), consistent with the idea that these sequences fold more reliably along trajectories leading to the targeted structures. Taken together, these results show that functional ribosensors can be generated through multi-state co-transcriptional folding design and provide a basis for investigating the mechanisms impacting device function.

### Kinetically-controlled binding and actuation

Three of the functional *p*-AF ribosensors were chosen for further study (Fig. 3a), including the devices with the largest predicted MFE path convergence (pAF4.5) (**Supplementary Table S2**), the largest and fastest rate of signal generation (pAF4.9) and the largest signal-to-noise ratio (pAF4.11) (**Supplementary Fig. S4**). Despite having identical aptamer sequences, the timer and toehold regions do not share sequence similarity (Fig. 3a, **Supplementary Table S4**), and have unique secondary structures predicted for the targeted ligand-binding (*s2*_*full*_) (Fig. 3a) and non-signaling OFF states (*s4*_*full*_) (**Supplementary Table S1**). Interestingly, the differences in the sequences and structures are reflected in the very different responses observed for each of the devices. pAF4.9 *(EC*_50_ = 360 ± 40 μM) and pAF4.11 (*EC*_*50*_ = 630 ± 180 μM) exhibited ligand-responsive outputs across concentrations spanning 3 orders of magnitude (12.5 μM to 12.5 mM) (Fig. 3a, **Supplementary Fig. S4**). And, both of these devices have very large ‘useful’ dynamic ranges^13^, such that the signal can be readily distinguished from noise across 40- and 60-fold differences in ligand concentration, respectively (Fig. 3a). By comparison, although pAF4.5 has the lowest *EC*_*50*_ (300 ± 50 μM), it also has a much narrower useful dynamic range (~12X).

**Figure 3.**
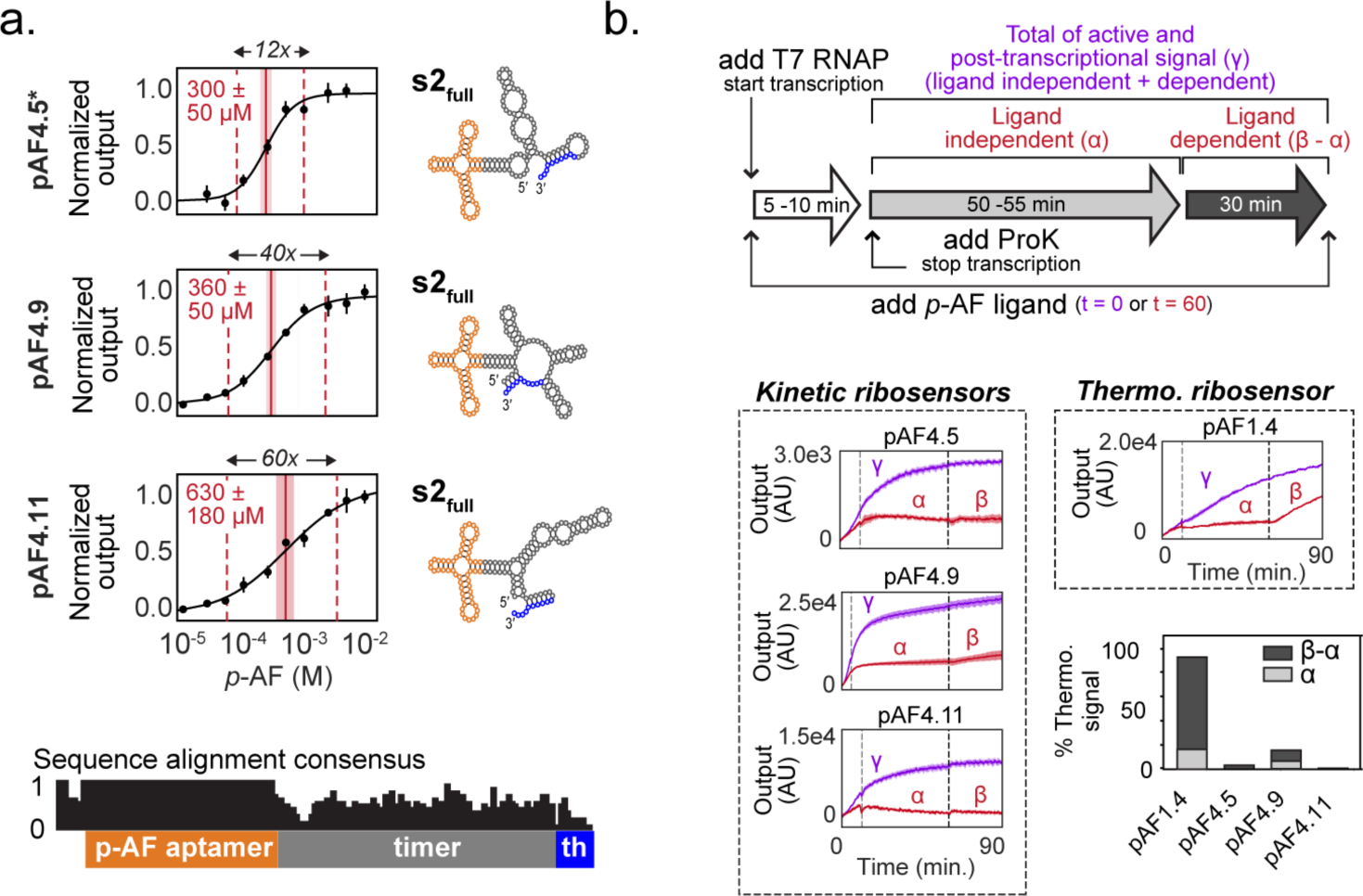
Ribosensors with kinetically-controlled binding and actuation. (a) The *EC*_*50*_s (red solid line), useful dynamic ranges (10% above the minimum and 10% below the maximum output; red dashed lines) and secondary structures of the final co-transcriptional states (s2_full_) are shown for each of three characterized ribosensors. (b) The aptamer sequence was held constant (orange), whereas there is little sequence similarity in the timer or toehold actuator (th, blue) domains. (b) The ligand-dependent (light gray, β-α) and ligand-independent (dark gray, α) post-transcriptional thermodynamically-controlled output of each ribosensor was isolated from the overall signal using a Proteinase K assay (top panel). The kinetic ribosensors pAF4.5, pAF4.9 and pAF4.11 display large (> 90%) kinetically-controlled, ligand-dependent responses. Ribosensor pAF1.4, designed to be under thermodynamic control, exhibits very little kinetically-controlled output (< 10%). Experiments were performed in triplicate. Standard deviations are indicated by shading.

To experimentally validate the kinetic control mechanism central to the design strategy, we developed an assay to measure how much of the observed ligand-dependent output was actually produced from the co-transcriptional, kinetically-controlled pathway (Fig. 1b). In the absence of ligand, ribosensors operating under kinetic control are designed to fold irreversibly into the non-signaling OFF state. The subsequent addition of ligand should not generate responses, except to the extent that ligand-dependent, thermodynamically-controlled transitions from the OFF state can occur (Fig. 1c). In this assay, we selectively halted transcription by degrading T7 RNA polymerase with Proteinase K (ProK) and compared the rates of co-transcriptional signal generation to the rates of post-transcriptional signal generation (Fig. 3b, **Supplementary Fig. S5**). In this way, we could isolate the contributions of the thermodynamically-controlled pathways from the overall signal and quantify the amount of kinetically-controlled ligand binding and actuation. Notably, we found that the thermodynamic pathways make only minor contributions to the overall signal generated by the pAF4.5 (3.2%), pAF4.9 (15.5%) and pAF4.11 (0.7%) devices (Fig. 3b, **Supplementary Figs. S6**, **S7**). For the pAF4.9 device, the background, constituting 6.2% of the overall signal, could be further separated (Fig. 3b), meaning greater than 90% of the ligand-dependent output from each of the three devices is under kinetic control (**Supplementary Fig. S7**).

In this architecture, the engineered binding window is the mechanistic feature designed to enable kinetic control. Accordingly, removing the co-transcriptional structure states and free energy barriers used to implement the binding window should minimize kinetic control. To test this hypothesis, we engineered pAF1.4 by eliminating a state (*s2*_*full*_) and a barrier (*B2*), intended to allow the RNA to fold directly into the non-signaling OFF state (*s4*_*full*_) (Fig. 1b, **Supplementary Table S1**). In addition, the free energy barriers that would otherwise separate the OFF state from the ligand-dependent, thermodynamically-controlled pathway (*B4* and *B5*) were minimized. As anticipated, the pAF1.4 device exhibits ligand-dependent function, but has almost entirely (92.2%) thermodynamic character, and is essentially devoid of kinetically-controlled, co-transcriptional ligand binding (Fig. 3b, **Supplementary Fig. S7**). Notably, the *EC*_*50*_ calculated for thermodynamic ribosensor pAF1.4 is significantly higher, at 2.8 ± 1.6 mM, than for any of the three kinetic ribosensors (**Supplementary Fig. S8**). This latter result supports the idea that kinetically-controlled responses can be significantly more sensitive to lower concentrations of ligand than thermodynamically-controlled responses^15^. These findings underscore the versatility of our design approach and indicate that the engineered binding window is essential for generating kinetic control within these devices.

### Cell-free devices for metabolic production biosensing

Many microbial pathways that are targets for metabolic engineering contain cytotoxic intermediates and enzymes whose overexpression disrupts the balance of cellular resources needed for growth and biosynthesis^18,30,31^. Thus, the process of identifying genetic designs to optimize flux through desired pathways for the production of medically- and industrially-relevant compounds typically requires screening large numbers of strain variants^32^. The current reliance on expensive, low-throughput and equipment-intensive analytical methods for screening^33,34^ constitutes a significant bottleneck in these efforts^32^. Because kinetic ribosensors can be transcribed *in situ* from inexpensive and shelf-stable components and rapidly generate fluorescent signals, they may be ideally-suited for metabolic production analysis and strain engineering^35^.

We first optimized a set of ribosensor assay conditions to allow for metabolite quantification using unfiltered cell supernatant taken directly from *E. coli* MG1655 and *E. coli* DH10b cultures (**Supplementary Fig. S9**). Although rapid RNA degradation has commonly hindered the application of aptamers as biosensors^12^, somewhat surprisingly, no RNA degradation was detected on the short timescales needed for these assays, showing that the ribosensors are stable enough for use in complex cell media (**Supplementary Fig. S9**). In the course of this optimization we noted that while high Mg^2+^ concentrations decrease the dynamic range of the outputs (**Supplementary Fig. S10**), neither ribosensor transcription nor TMSD actuation are significantly impacted by the presence of 50% MOPS EZ-Rich microbial cell supernatant (**Supplementary Figs. S9, S10**).

Next, we evaluated the pAF4.9 and pAF4.11 device functions in simulated production assay conditions with the addition of exogenous *p*-AF to spent *E. coli* media (Fig. 4a, **Supplementary Fig. S11**). While the *EC*_*50*_s (820 ± 160 μM and 500 ± 170 μM, respectively) are slightly higher in 50% MOPS EZ-Rich media when compared to the standard assay conditions, the useful dynamic ranges of both devices nonetheless encompass concentration ranges with immediate relevance for metabolic engineering^19^. pAF4.9, in particular, rapidly generates a high level of output signal (**Supplementary Fig. S4**) and has a high signal to noise ratio, suitable for high-throughput screening.

**Figure 4.**
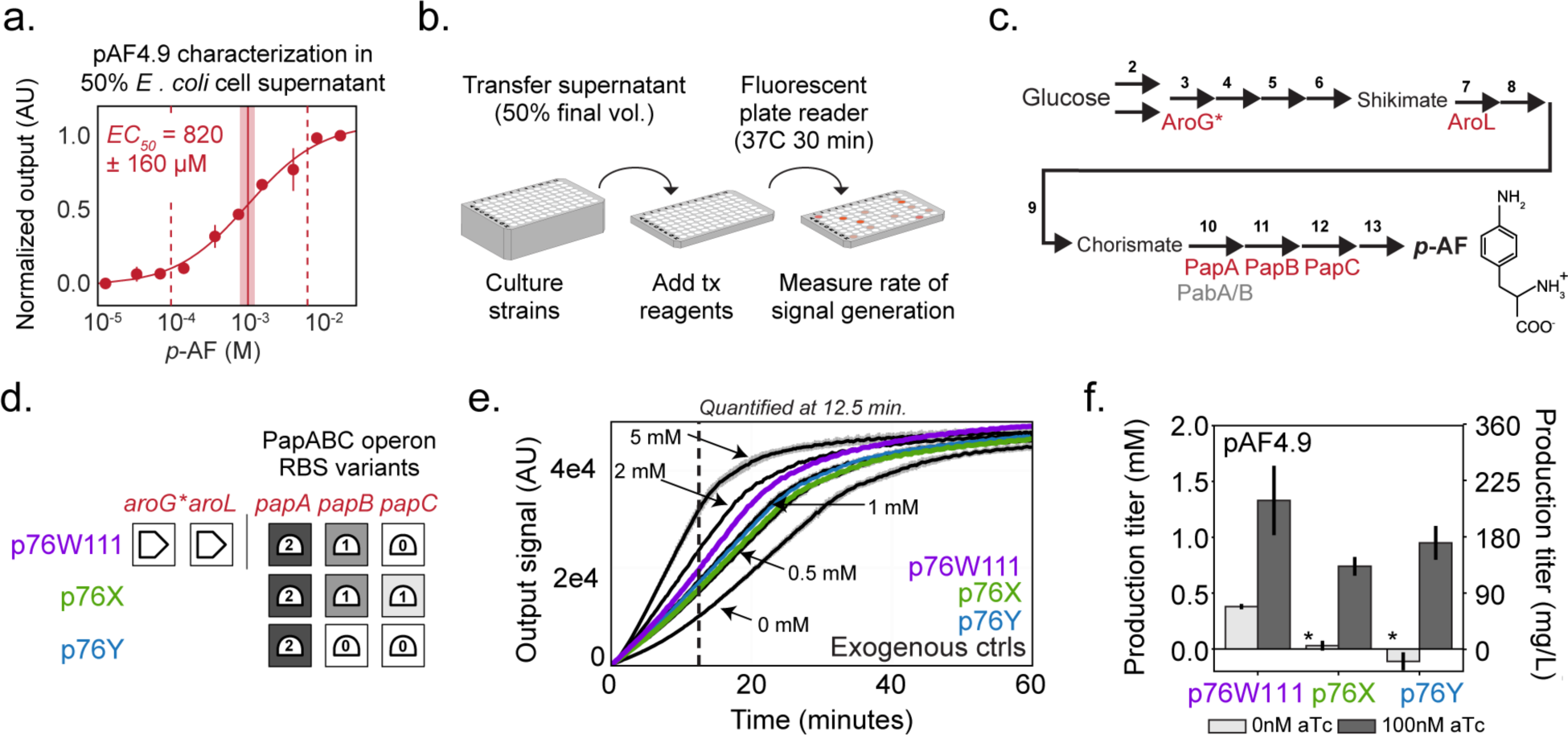
Ribosensors for screening production phenotypes. (a) In 50% *E. coli* cell supernatant pAF4.9 has a useful dynamic range (red dashed lines) spanning ~50-fold differences in *p*-AF concentration. (b) A simple workflow for metabolic production analysis was devised. (c) Engineered *p*-AF production pathway with heterologous genes shown in red and endogenous genes expected to impact pathway flux shown in gray. (d) Metabolic production analysis was performed on three engineered *p*-AF production pathway variants. (e) Fluorescence outputs generated by the pAF4.9 ribosensor are plotted as a function of time for each of the three induced pathway variants (purple, green, blue) and exogenously *p*-AF controls (black). (f) Metabolic flux through the engineered pathway was quantified by comparing production titers to a standard curve (Supplementary Fig. S11). * indicates that the signal was below the range of the standard curve. Experiments were performed in triplicate. Error bars and shading indicate standard deviation.

Finally, to demonstrate that ribosensors can be employed for production analysis (Fig. 4b), the pAF4.9 device was used to quantify metabolic outputs from a set of *E. coli* variants engineered to direct flux through the *p*-AF biosynthesis pathway (Figs. 4c, 4d, **Supplementary Fig. S1**). A simple assay workflow was devised: media samples from cell cultures are added directly to ribosensor components without any filtration or additional preparation. Final reaction volumes are 50% cell culture supernatant, requiring only minimal, 2-fold dilution of the analyte.

Ribosensor outputs are then measured in real-time over the course of the next 10-30 minutes (Fig. 4e). Clear differences in ribosensor output are seen when heterologous gene expression is induced compared to the no-inducer controls (Fig. 4f). And further, by comparison to pAF4.9 outputs generated in response to exogenous *p*-AF, metabolic production titers from the engineered strain variants were easily quantified (p76X: 0.75 ± 0.08 mM, p76Y: 0.95 ± 0.15 mM, p76W111: 1.33 ± 0.31 mM) (Fig. 4f, **Supplementary Fig. S12**).

### Enhancing ligand sensitivity and selectivity through kinetic control

We next turned to the questions of whether, and how, kinetic control can be implemented to provide unique capabilities for enhancing ribosensor sensitivity and selectivity that cannot be obtained through thermodynamic control. For thermodynamically-controlled RNAs, assembling an aptamer into a more complex structure involving additional sequence elements will necessarily create competition between the equilibrium ON and OFF states. Consequently, the sensitivity of thermodynamic devices is inherently limited such that concentrations of ligand much higher than the aptamer-ligand dissociation constant (*K*_*d*_) are required to generate a response^36^. In contrast, for kinetically-controlled devices, the sensitivity of the response is related to the size of the binding window, or amount of time the aptamer domain can associate with the ligand. Increasing the duration of the binding window should, all else being equal, increase the sensitivity of the device to lower concentrations of ligand^5^. By the same token, when aptamer association rates for a set of cognate ligands are distinct, kinetic control can provide molecular discrimination even though differences in the equilibrium aptamer-ligand binding constants may be very small^6^.

Formally, the sensitivity of the ribosensor response is dictated by the ligand association rate (*k*_*on*_), an intrinsic property for an aptamer-ligand pair, and the time of the binding window (*t*_*bind*_) (Fig. 1b), a parameter that can be tuned. As a proof-of-concept, we reduced the total concentration of ribonucleotide substrates (rNTPs) from 4 mM to 0.4 mM in a *p*-AF titration assay with the pAF4.9 ribosensor. Reducing the concentration of rNTPs should decrease the T7 RNA polymerase elongation rate, extend the binding window and increase the sensitivity of the device to lower concentrations of *p*-AF^2,37,38^. Consistent with expectation, a two-fold increase in the sensitivity of the pAF4.9 device was observed, with the *EC*_*50*_ dropping from 360 ± 40 μM to 180 ± 50 μM upon reducing the rNTP concentration (Fig. 5).

**Figure 5.**
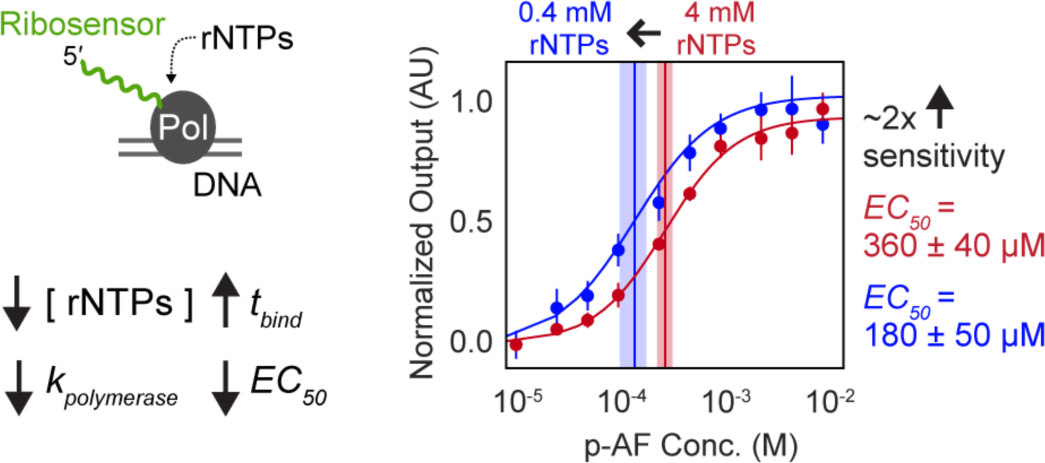
Kinetic control of ligand sensitivity. Predictable increases in the sensitivity of the pAF4.9 ribosensor were obtained by reducing the concentration of rNTP substrate. Changing the concentration of rNTP substrate from 4 mM to 0.4 mM was expected to reduce the rate of polymerase elongation, increase the duration of the binding window (*t*_*bind*_) and shift the response of the device to lower concentrations of *p*-AF. Vertical lines correspond to the calculated *EC*_*50*_s. Experiments were performed in triplicate and data are reported with standard deviation (error bars and shading)

To show that kinetic control can modulate the selectivity of the ribosensor response, we applied the design rules developed to engineer the *p*-AF devices to assemble an aptamer promiscuous for aromatic amino acid binding into kinetically-controlled ribosensors (**Supplementary Fig. S13**). We then measured the responses of one of the resulting devices, Tyr1.5 (Fig. 6a, **Supplementary Fig. S14**), to a panel of cognate ligands. Interestingly, even though the parental Tyr1b aptamer has the same measured *K*_*d*_ for L-Trp (22 ± 1 μM), L-Tyr (23 ± 2 μM), and L-DOPA (22 ± 3 μM)^20,21^, the Tyr1.5 ribosensor has strikingly different responses to this panel of ligands (Fig. 6b, **Supplementary Fig. S14**). The Tyr1.5 ribosensor response to 1 mM L-Trp is 2-times as large as the response to 1 mM L-Tyr and, remarkably, 45-times as large as the response to 1 mM L-DOPA (Fig. 6c). Furthermore, even though the parental aptamer *K*_*d*_ (183 ± 26 μM) is 9-times higher for L-Phe than for L-DOPA, the Tyr1.5 ribosensor nonetheless exhibits a larger response to L-Phe than to L-DOPA (Fig. 6c, **Supplementary Fig. S14**). These data show that kinetic control can be employed to generate ligand selectivity, without re-engineering the parental RNA aptamer^39^, that is beyond what would be possible through differences in the thermodynamics of molecular recognition.

**Figure 6.**
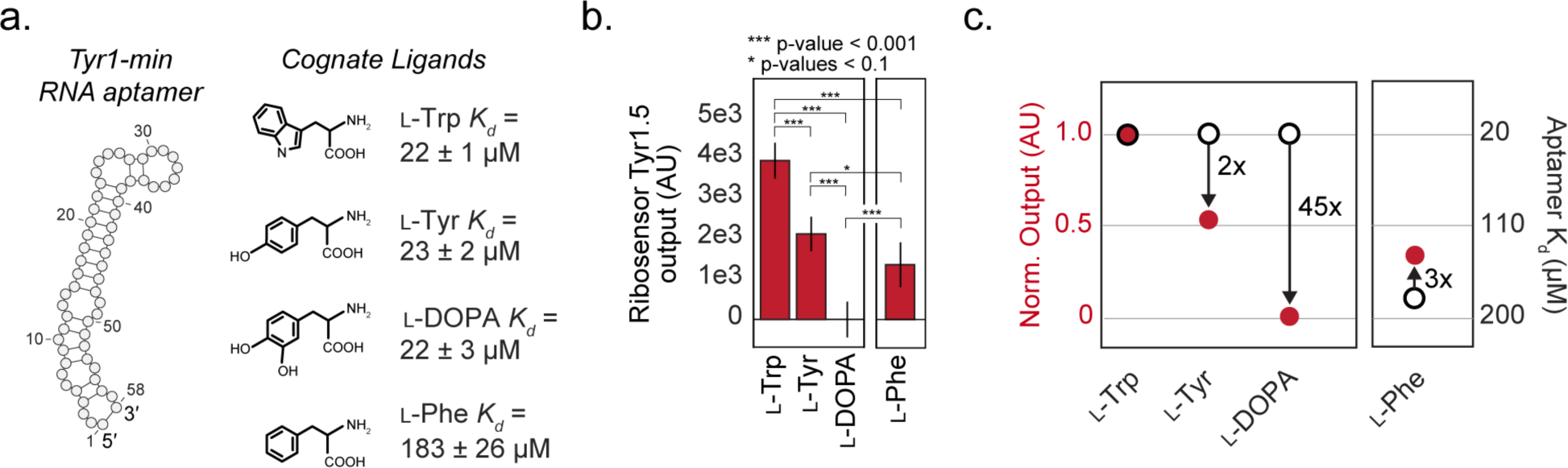
Kinetic control of ligand selectivity. (a) The Tyr1 aptamer is promiscuous for aromatic amino acid binding. (b) The kinetically-controlled Tyr1.5 ribosensor was engineered from the Tyr1 aptamer by applying the design rules developed for the *p*-AF ribosensors. The bar chart shows the responses of the Tyr1.5 ribosensor to 1 mM of the designated cognate ligand. (c) Tyr1.5 ribosensor outputs with the cognate ligands (red circles, left axis) are normalized to the response generated with 1 mM L-Trp and plotted alongside the *K*_*d*_ of the parental aptamer (open circles, right axis). This representation illustrates the differences in ligand selectivity obtained with kinetic control against what would be expected through the thermodynamics of aptamer molecular recognition (arrows). Experiments were performed in triplicate and data are reported with standard deviation.

## DISCUSSION

The role of kinetic control mechanisms in creating biological activities that are both sensitive to, and selective for, specific molecular interactions has long been recognized^10^. Next-generation RNA aptamer biosensors have been successfully created through *in vitro* selection^40^, directed evolution^7^, activity-based screening^41,42^, two-state RNA folding design^8^, and thermodynamic optimization^43^. However, none of these state-of-the-art methods routinely generate RNA devices operating under kinetic control^7,8^, and an entirely different strategy was required to access that class of mechanisms. Here, we successfully assembled two different *in vitro* selected aptamers into kinetically-controlled ribosensors through multi-state co-transcriptional RNA folding design^17,20^. There was excellent correspondence across the set of 29 candidate *p*-AF ribosensors between device function, and both the presence of the targeted secondary structure states and degree of convergence along the predicted co-transcriptional folding trajectories (Fig. 2c). Consistent with the design intent, >90% of the output generated by each of three characterized devices originated from the ligand-dependent, kinetically-controlled pathway (Fig. 3b). Removing the secondary structures and free energy barriers framing the ligand binding window all but eliminated kinetic control in the pAF1.4 device (Fig. 3), highlighting the role of the binding window in conveying co-transcriptional, kinetically-controlled responses. Thus, our novel approach constitutes a significant conceptual and practical advance toward the reliable design of kinetically-controlled RNA aptamer devices.

Engineered biosensors producing detectable signals are in high demand as replacements for expensive and equipment-intensive analytical methods^33,34^. The engineered ribosensors are transcribed *in situ* and can be used to quantify production titers directly from engineered *E. coli* with a very simple workflow. Whereas traditional HPLC-based methods for quantifying metabolic production may take 10-30 minutes to analyze a single sample, an entire 96- or 384-well plate of samples can be analyzed using ribosensors in the same amount of time (e.g. 12 minute assays for production analysis with the pAF4.9 device, Fig. 4). Because the sensitivity of thermodynamically-controlled aptamer devices, including biosensors commonly engineered with the fluorogenic Spinach-family of aptamers, are fundamentally constrained by the equilibrium competition between the ON and OFF states^40,41,43^, they will always require concentrations significantly higher than the aptamer-ligand dissociation constant (*K*_*d*_) for a response^13,36^. In this regard, it is very promising that the *EC*_*50*_s of the three kinetically-controlled *p*-AF ribosensors are both consistent with values predicted from the underlying design parameters (**Supplementary Fig. S15**), and are nearly 5-10 times lower than the *EC*_*50*_ of the thermodynamic pAF1.4 ribosensor (300-680 μM versus 3 mM, respectively) (Fig. 3a, **Supplementary Fig. S8**). As a practical consequence, this difference in sensitivity permits *p*-AF concentrations as low as 70 μM (12.5 mg/L) to be quantified using the kinetically-controlled ribosensors, well below the levels detectable by the thermodynamic device (220 μM) (Fig. 3a, **Supplementary Fig. S8**). The application of the ribosensor design approach provides a new route to extensible biosensors with immediate utility for rapidly analyzing combinatorial genetic design spaces and optimizing high-value chemical production^18,32^. For instance, the chorismate biosynthesis superpathway, from which *p*-AF is produced, is the origin of a vast array of medically- and industrially-relevant aromatic compounds (**Supplementary Fig. S1**)^44,45^. Cognate RNA aptamers for a number of high value aromatics already exist and additional aptamers for intermediates within, or outside, this pathway can be evolved with *in vitro* selection^40,46–49^. Notably, many of the same properties useful for metabolic production analysis, including inexpensive (<$0.40/reaction) and freeze-dryable components, and a rapid rate of signal generation, make kinetic ribosensors well-suited as point-of-care diagnostics. By assembling ribosensors from aptamers for small molecule or protein biomarkers, plate- or paper-based assays could be developed for applications in medicine, biodefense or global health^50,51^.

Because ribosensors employ toehold-mediated strand-displacement actuators, multiplexed analyses could be performed by transcribing multiple ribosensors in the same reaction and then quantifying the outputs with spectroscopically-distinct fluorophores, nucleic acid probe hybridization, or NGS sequencing^16^. By comparison, the potential for multiplexed analysis with aptamer biosensors employing fluorogenic aptamers as actuators will continue to be limited unless additional aptamer-fluorophore pairs are created^52^.

The ability to design kinetically-controlled RNA aptamer devices creates unique capabilities for programming the sensitivity of the response to lower concentrations of ligand or selectively discriminating among cognate ligands with similar equilibrium binding constants^12,35^. Predictable, two-fold enhancements in *p*-AF ribosensor concentration sensitivity were easily obtained by slowing the RNA polymerase and increasing the amount of time needed to transcribe the timer domain (Fig. 5). Looking ahead, much larger increases in the duration of the binding window, and thus ligand sensitivity, could be instantiated with longer timer sequences, transcriptional pause sites^5^, or even slowly-resolving post-transcriptional kinetic traps, theoretically allowing the *EC*_*50*_ of the assembled devices to approach the equilibrium *K*_*d*_ of the aptamer alone (**Supplementary Fig. S15**). Remarkably, the Tyr1.5 kinetic ribosensor generated 45-fold differences in the selectivity of the response to L-Phe compared to L-DOPA, even though the parental aptamer exhibits the same *K*_*d*_ for both targets (Fig. 6)^20,21^. Developing the capacity to specify the exact duration of the binding window, through a combination of biochemical tuning and RNA folding design, may enable ribosensor responses to be precisely matched to targeted levels of sensitivity and selectivity. Using this work as a starting point, multi-state co-transcriptional RNA folding design may eventually permit the fully-rational engineering of RNA aptamer devices for a wide range of architectures and applications in biosensing or genetic control.

## METHODS

### Materials and Methods

#### Reagents and oligonucleotides

All DNA oligonucleotides were purchased from Integrated DNA Technologies (Coralville, IA). Ribosensor templates were purchased as Ultramers, modified oligonucleotides for DNA strand displacement gates were purified by HPLC by IDT, all other oligonucleotides were used without purification, unless otherwise stated. 4-amino-phenylalanine was purchased from Santa Cruz Biotechnology (Dallas, TX) and made fresh for each experiment. L-tyrosine, L-phenylalanine, 3,4-Dihydroxy-L-phenylalanine (L-DOPA), L-tryptophan, were purchased from Sigma-Aldrich (St Louis, MO) and stocks made fresh for each experiment (10 mM in 10 mM HCl).

Anhydrotetracycline hydrochloride (aTc), ammonium persulfate, trifluoroacetic acid (TFA), spermidine, potassium glutamate, dithiothreitol (DTT), and potassium chloride were also purchased from Sigma-Aldrich (St Louis, MO). T7 RNA polymerase and ribonucleotides were purchased from New England Biolabs Inc. (Ipswich, MA). 2x DreamTaq Master mix and carbenicillin sodium salt were purchased from Thermo Fisher Scientific (Waltham, MA).

Proteinase K lyophilized powder and *E. coli* DH10b chemically competent cells were purchased from Invitrogen (Carlsbad, CA). Nucleic acid silica membrane spin columns were purchased from Epoch (Missouri City, TX) and were used with commercially available P1, P2, N3, PB, PE, and EB buffers from Qiagen (Germantown, MD). LB Miller media and MOPS EZ-Rich media were purchased from Teknova (Hollister CA). 30% acrylamide/bis solution, 19:1 was purchased from BioRad (Hercules, CA).

#### p-AF ribosensor design

*p-*AF ribosensors were engineered from the *p*AF4z1d3 RNA aptamer^17^. We designed initial *in silico* libraries using a ribosensor architecture which contained a 5′ RNA aptamer followed by a 65 nucleotide timer and ending with a 3′, 10 nucleotide toehold actuator domain (148 nucleotides total in length). The architecture was used to define three full-length secondary structure states *s2*_*full*_, *s3*_*full*_ and *s4*_*full*_ (**Supplementary Fig. S2**). The aptamer domain sequence was held constant with the remainder of the sequence allowed to vary (**Supplementary Table S4**). These structure states were used as target secondary structures for the inverse folding, multi-objective genetic algorithm, MODENA (ver0.0.67)(**Supplementary Fig. S2**)^26^; house-written software was employed for set-up and analysis on a computational cluster. Because MODENA utilizes an inverse folding approach, 2 objective functions (OFs) were defined to screen the relative free energies of each target state for a given generated sequence (ΔG s4_full_ = MFE, and ΔG s3_full_ ≅ ΔG s2_full_) (**Supplementary Fig. S2**). In addition, a third OF was specified to maximize sequence and structural constraint similarities between the input target and output structure states (**Supplementary Fig. S2**). Since the MODENA algorithm returns Pareto optimal solutions that may not necessarily meet all of specified sequence and structure constraints, the outputs were re-screened using the input structures and sequences as constraints. Sequences were judged to have met the screening criteria when the secondary structure free energies of the targeted structure states fell into the following rank order: *s3*_*full*_ > *s2*_*full*_ > *s4*_*full*_

(**Supplementary Table S1**). Thus, the *in silico* ribosensor pools generated using this method (**Supplementary Fig. S2**) contained pre-screened sequences with the potential to fold into the targeted full-length structure states *s2*_*full*_, *s3*_*full*_ and *s4*_*full*_ and satisfied the targeted multi-state free energy relationships.

Next, the accessibility of the structure states along the predicted co-transcriptional folding trajectories was evaluated for all candidate ribosensor sequences using our house-written *MFEpath* algorithm (**Supplementary Fig. S3**). The MFE threshold was set to 6.4 kcal/mol defined by the upper limit of T7 RNAP transcription rates (200 nts/sec)^25,27,53^. The free energy relationships among the targeted structure states (i.e., *s1*_*co-txn*_, *s2*_*full*_ and *s4*_*full*_, as shown in **Supplementary Table S1**) were evaluated when those states were found to be present along the *MFEpath*-predicted folding pathways (**Supplementary Table S2**). Finally, direct folding pathways were analyzed to calculate the rates of transition from the final, *MFEpath*-predicted, post-transcriptional structure (*s2*_*full*_) to either the ligand-bound actuation state (*s3*_*full*_) or the ligand-unbound OFF state (*s4*_*full*_) (**Supplementary Table S3**). The *FindPath* binary from the Vienna RNA package (ver2.1.9) was employed for this purpose^29^, using custom software for set-up and analysis on a local computer. Sequences were judged to have met the barrier height design criteria when the following rank order was achieved: *B3* and *B4* > *B1* > *B2* (**Supplementary Table S3**).

Note that a subset of sequences were also evaluated using stochastic co-transcriptional folding simulations to evaluate the accessibility of the co-transcriptional structure states (*s1*_*co-txn*_ and *s2*_*full*_). House-written software built on the *Kinefold* binary was employed for set-up and analysis^54,55^. Sequences were judged to be successful when the following criteria were met: 1) the RNA aptamer structure appeared at a high frequency (>50% of simulated runs) along the predicted folding trajectory, while the functional toehold actuator did not appear, and 2) the *s4*_*full*_ state was present at the completion of transcription. Subsequent analysis, however, determined that this computationally-intensive *Kinefold*-based co-transcriptional screening step is not needed to obtain functional devices as long as the *MFEpath*-derived structure state and barrier height design criteria are met (as above).

### Thermodynamic p-AF ribosensor design

The thermodynamic *p*-AF ribosensor was engineered using the same pAF4z1d3 RNA aptamer as the kinetic ribosensors^17^. The timer length was shortened to 29 nucleotides (nt) to decrease *t*_*bind*_ and the toehold region was kept at 10 nt (**Supplementary Table S4**). A successful thermodynamic sequence was judged to have no *s2*_*full*_ state or barrier *B2* along the folding trajectory, contrary to the kinetic counterparts described above (**Supplementary Tables S1**, **S3**). In addition, sequences were judged to have met the barrier height design criteria when the following rank order was achieved: *B3* > *B4* and *B5* (**Supplementary Table S3**).

### Aromatic amino acid ribosensor design

To engineer aromatic amino acid ribosensors, the Tyr1-min sequence and the MFE secondary structure predicted using *RNAfold* (Vienna RNA package) were substituted for the sequence and structure of the pAF4z1d3 aptamer and processed using the same computational pipeline and design criteria as above, without modification (**Supplementary Fig. S2, Supplementary Table S4)**^20^. Five candidate ribosensors were generated that met the design criteria for 1) the full-length sequence, secondary structure free energies, and secondary structure states 2) the *MFEpath*-derived co-transcriptional structure states and barrier-heights, and 3) the post-transcriptional barrier heights. One of the resulting ribosensors, Tyr1.5, exhibited ligand-dependent outputs with 1 mM L-Tyrosine and was used for subsequent analysis (**Supplementary Fig. S13, Supplementary Table S4**).

### DNA gate preparation

For 4-stranded X-Probe gate preparation^16^, 10 μM each DNA strand (**Supplementary Table S5**) were combined in 10 mM Tris with 10 mM MgCl_2_. Samples were heated to 95 C for 1 minute and cooled −1 C / minute until 25 C was reached. Gates were purified using a 10% native polyacrylamide gel run at 3 W for 30 minutes. Gate bands were UV-shadowed and excised from the gel. The gate was eluted from the gel in 300 μL of 300 mM KCl by overnight shaking, then recovered by ethanol precipitation. The gates were suspended in 30 μL of 10 mM Tris and the concentration was quantified by absorbance at 260 nm on a Nanodrop spectrophotometer. For 2-stranded gate preparation, 50 mM each strand was used with the remainder of the protocol unchanged.

### Ribosensor template preparation

DNA templates (**Supplementary Table S4**) for *in vitro* transcriptions were prepared through PCR amplification using DreamTaq 2x Master Mix, 1 nM template, and 1 μM each primer, where the forward primer was used to add the T7 RNA polymerase promoter sequence (**Supplementary Table S4**). Primers were design to have a melting temperature between 52 – 55 C. Templates were subsequently diluted (1000x) and re-amplified with 1 μM primers (T7 Promoter specific forward to ensure that the 5′ end of the templates were clean, and the same device specific reverse primer used above). PCR products were purified using silica membrane spin columns and template concentration was quantified at 260 nm on a Nanodrop spectrophotometer.

### General procedure for in vitro ribosensor fluorescence assays

20 μL reactions with final concentrations of 40mM Tris pH 8, 6 mM MgCl_2_, 2 mM spermidine, 10 mM DTT, 150 mM potassium glutamate, 1 mM each rNTP, 50 U T7 RNA polymerase, 30 ng ribosensor template, 5 ng control template, 100 nM X-Probe ROX labeled ribosensor gate, 100 nM 2-stranded FAM labeled control gate, were combined on ice in white PCR tubes with optically clean caps. For *p*-AF ribosensors, ligand concentration ranged from 12.5 μM to 12.5 mM, and for the aromatic amino acid sensor the cognate ligands (L-Trp, L-Tyr, L-Phe, and L-DOPA) were assayed at 1 mM. Fluorescence was measured at 37 C every 15 sec. in an Mx3005p qPCR system (Agilent) using the ROX (588 nm / 608 nm) and FAM (492 nm / 516 nm) filter sets. For titration analysis, the data was fit to a Hill function in Python with the *EC*_*50*_ reported with the standard deviation of at least 3 replicates.

### Proteinase K assay

20 μL reactions with final concentrations of 40mM Tris pH 8, 6 mM MgCl_2_, 2 mM spermidine, 10 mM DTT, 150 mM K-glutamate, 1 mM each rNTP, 50 U T7 RNA polymerase, 30 ng Ribosensor template, 5 ng control template, 100 nM X-Probe ROX labeled Ribosensor gate, 100 nM 2-stranded FAM labeled control gate, were combined on ice in white PCR tubes with optically clean caps. Ligand concentrations are indicated in the Figures, but were either 12.5 mM or 0 mM. Reactions were run at 37 C in the Mx3005p qPCR system. Transcription was halted between 5 and 10 minutes by the addition of 2U Proteinase K. Fluorescence was measured at 37 C every 15 sec. for 1 hour. 12.5 mM ligand was then added and fluorescence was measured at 37 C every 15 sec. for an additional 30 minutes.

### Plasmid construction

All plasmids were constructed using circular polymerase extension cloning (CPEC). All cloning-related transformations were performed with *E. coli* DH10b cells. Plasmid backbones were derived from pBbE2a-RFP (Addgene #35322). The 76W/X/Y strains are RBS variants of the same expression cassette. The Tn10-derived ptet promoter controls expression of a three-gene operon consisting of PapA, PapB, and PapC. The backbone, containing the tet promoter, BBa_B0015 terminator, ColE1 origin of replication, and ampicillin resistance marker was taken from the pBbE2A plasmid^56^. RBS sequences were taken from Mehl, et al. 2003^57^ or generated using the RBS Calculator v2.0^58^. PapB and PapC are codon-optimized sequences of the *S. venezuelae* strain (ATCC10712) cited in Mehl, et al. 2003^57^. The *papA* gene was generated using sequence data from constructs in Carothers et al. 2011^9^, originally obtained as a gift from Prof. Chris Anderson, and was also codon optimized for expression in *E.coli.* Genes were synthesized by GeneArt as Strings DNA Fragments (ThermoFisher).

p111 is an expression cassette with two genes in the *E. coli* aromatic amino acid synthesis pathway. These two genes are aroG* (feedback-resistant mutant) and aroL. Both are natively regulated by the transcriptional regulator tyrR, so it is expected that heterologous genes not subject to tyrR regulation could increase flux toward chorismate, the starting point for making *p*-AF. The Tn10-derived ptet promoter controls expression of aroG* and aroL as an operon. The backbone, containing the tet promoter, BBa_B0015 terminator, ColE1 origin of replication, and ampicillin resistance marker was taken from the pBbE2A plasmid^56^. The sequences for aroG* and aroL were amplified from pS4 and pY3, respectively, from Juminaga, et al. 2012^45^. The RBS for aroG* is the same as that of pJTS76Y, and the RBS for aroL was taken from its RBS in pY3.

### Production assay

*E. coli* DH10b chemically competent cells were transformed with plasmid(s) and plated on LB agar plates with carbenicillin antibiotic. Five distinct colonies were picked to inoculate 1 mL of LB Miller media with 100 μg/mL carbenicillin sodium salt in 14 mL polypropylene culture tubes. Cultures are grown at 37 C and shaken at 200 rpm overnight. 20 μL of overnight culture is used to inoculate 2 mL of MOPS EZ-Rich media and 100 μg/mL carbenicillin. Gene expression was induced with 100 nM aTc when OD600 reached 0.3-0.4. Cultures were then grown at 37 C with 200 rpm shaking for 24 hours.

### Ribosensor measurements from production strains

Production cultures were centrifuged at 4000 g for 15 minutes. 10 μL of supernatant were then added to 20 μL ribosensor assay reactions, with final concentrations of 50% (v/v) cell culture supernatant, 6 mM MgCl_2_, 10 mM DTT, 150 mM K-glutamate, 1 mM each rNTP, 50 U T7 RNA polymerase, 30 ng ribosensor template (pAF4.9), 5 ng control template, 100 nM X-Probe ROX labeled Ribosensor gate and 100 nM 2-stranded FAM labeled control gate. Reaction components were combined on ice in white PCR tubes with optically clean caps. Fluorescence was measured at 37 C every 15 sec. in an Mx3005p qPCR system (Agilent) using the ROX (588 nm / 608 nm) and FAM (492 nm / 516 nm) filter sets.

### rNTP titration for sensitivity tuning

20 μL reactions were assayed with final concentrations of 40mM Tris pH 8, 6 mM MgCl_2_, 2 mM spermidine, 10 mM DTT, 150 mM K-glutamate, 1 or 0.1 mM each rNTP, 50 U T7 RNA polymerase, 30 ng ribosensor template, 5 ng control template, 100 nM X-Probe ROX labeled Ribosensor gate and100 nM 2-stranded FAM labeled control gate. Reaction components were combined on ice in white PCR tubes with optically clean caps. *p*-AF ligand concentrations ranged from 12.5 μM to 12.5 mM. Fluorescence was measured at 37 C every 15 sec. in an Mx3005p qPCR system (Agilent) using the ROX (588 nm / 608 nm) and FAM (492 nm / 516 nm) filter sets.

## Acknowledgements

We thank Prof. Georg Seelig, Randolph Lopez and Sifang Chen for advice on Toehold-Mediated Strand Displacement (TMSD) and Prof. David Zhang for pre-publication X-Probe access. We thank Chuhern Hwang for sharing aptamer binding data and Jason Stevens for sharing engineered *p*-AF pathway strains. Thanks to Profs. Georg Seelig and Jesse Zalatan for feedback on the manuscript. This work was supported in part by funds from NSF Award MCB 1517052 and a University of Washington Presidential Innovation Award. J.M.C. was a fellow of the Alfred P. Sloan Foundation.

## Author contributions

C.R.B. and J.M.C. conceived of the project, designed experiments and wrote the manuscript with contributions from all authors. C.R.B. performed data collection. D.S-Y. developed *MFEpath*.

